# Broad sialic acid usage amongst species D human adenovirus

**DOI:** 10.1101/2023.03.21.533702

**Authors:** Rosie M. Mundy, Alexander T. Baker, Emily A. Bates, Tabitha G. Cunliffe, Alicia Teijeira-Crespo, Elise Moses, Pierre J. Rizkallah, Alan L. Parker

## Abstract

Human adenoviruses (HAdV) are widespread pathogens causing infections of the respiratory and gastrointestinal tracts, genitourinary system and the eye. Species D (HAdV-D) are the most diverse species and cause both gastrointestinal tract infections and epidemic keratoconjunctivitis (EKC). Despite being significant pathogens, HAdV-D are understudied and knowledge around basic mechanisms of cell infection is lacking. Sialic acid (SA) usage has been proposed as a major mechanism of cell infection for EKC causing HAdV-D. Here, we provide apo state crystal structures for fiber knob proteins of 7 previously undetermined HAdV-D, and provide crystal structures of HAdV-D25, HAdV-D29 and HAdV-D53 knob proteins bound to SA. Biologically, we demonstrate that removal of cell surface SA reduced infectivity of HAdV-C5 vectors pseudotyped with HAdV-D fiber knob proteins, whilst engagement of the classical HAdV receptor, CAR was variable. Together, these data indicate an important role for SA engagement in the tropism of many HAdV-D and may facilitate the development of suitable antivirals to control EKC outbreaks.

## Introduction

Human adenoviruses (HAdV) are classified phylogenetically into seven species, A-G, based classically on testing for neutralizing activity by serological analysis (*1, 2*). They are highly diverse, and have become of great interest for their use as therapeutic vectors and vaccines. Adenoviruses are non-enveloped viruses and have a double stranded DNA genome of around 35kb, which lends itself readily to genetic manipulations (*3*). These manipulations allow them to be tailored into safe and effective delivery platforms for gene therapy and oncolytic payloads which, coupled with their good safety profile, make them attractive targets for therapeutic development.

Adenoviruses have a highly conserved structure (*4*). Their icosahedral capsid is made up of three major proteins: the hexon, penton and fiber, in addition to other minor and cement proteins. Each protein forms key interactions to promote cell infection, with the terminal domain of the fiber, the fiber knob protein, interacting with the viruses’ primary cell entry receptor.

Therapeutic development has historically focused significantly on the archetype species C adenovirus type 5 (HAdV-C5). However, the use of this virus faces many hurdles including high pre-existing seroprevalence rates in some populations, as well as undesirable interactions with blood components such as blood clotting factors (*5, 6*), which may reduce efficacy and promote off target toxicities of platforms based on HAdV-C5 (*7–9*). Species D adenoviruses (HAdV-D) represent alluring alternatives, with generally lower rates of pre-existing immunity in Western populations, however they are severely understudied when compared to others, such as the species C adenoviruses.

Species D adenoviruses generally cause ocular and gastrointestinal infections, including HAdV-D8, which recently caused an outbreak of conjunctivitis in Leicester, UK (*10*). They can also cause epidemic keratoconjunvtivitis (EKC), which is endemic to Japan and causes a strain on the healthcare system (*11*). The lack of understanding on species D viruses’ basic mechanisms of cellular infection prevents better treatment of these diseases, and clarification would allow generation of more effective therapeutics.

Previous work has demonstrated that species D adenovirus 26 (HAdV-D26) fiber knob protein does not engage classical proteinaceous receptor CD46, and only weakly binds Coxsackie and Adenovirus Receptor (CAR), where its engagement is sterically hindered by an extended DG loop, blocking this interface. Instead, the HAdV-D26 fiber knob protein engaged sialic acid with high affinity as a cell entry receptor (*12, 13*). HAdV-D26 has been used extensively clinically as vaccines against Ebola (*14*), HIV (*15*) and most recently SARS-CoV-2 (*16*). Interactions with sialic acid have been described with other species D adenoviruses including adenovirus 19p (HAdV-D19p), 37 (HAdV-D37) and 64 (HAdV-D64) (*17–19*). Sialic acid is also known to be commonly used in other virus infection mechanisms such as influenza and is the terminal residue of complex glycan chains on a large proportion of vertebrate cell surfaces, including humans (*20*). All of the aforementioned viruses bind sialic acid in an apical binding pocket, despite HAdV-D26 fiber knob protein having a different electrostatic profile and low sequence similarity to the others (*12*), suggesting the potential for widespread usage of the receptor across the species D adenoviruses. We assessed the ability for HAdV-D to engage sialic acid and CAR as cell entry receptors via an integrative structural biology workflow, suggesting a species wide dual tropism,

## Methods

### Generation and purification of recombinant fiber knob proteins

pQE expression plasmids (Qiagen, Manchester, UK) containing DNA encoding species D fiber knob proteins were produced by TWIST Biosciences (San Francisco, USA). The vectors were transformed by heat shock into competent SG13009 *Escherichia coli (E.coli*) bacteria containing a pREP4-4 plasmid. Heat shock transformation was performed by heating the SG13009 *E.coli* and pQE-30 vectors at 42°C in a waterbath for 30 seconds and cooling on ice for 2 minutes.

250μL of room temperature SOC media (Super Optimal Broth with added glucose) was added and bacteria recovered in a shaking incubator for 1 hour at 37°C. Transformed bacteria were spread on LB-agar plates with 100μg/mL Kanamycin and 50μg/mL Ampicillin and grown overnight at 37°C. Transformed colonies were confirmed by sequencing. SG13009 colonies containing the appropriate species D pQE fiber knob protein expression vectors were cultured to an optical density of 0.6 in Terrific Broth (TB, Sigma-Aldrich, Gillingham, UK), harvested by centrifugation and purified by nickel immobilised metal affinity chromatography (IMAC) and size exclusion chromatography (SEC) as described previously (*12*).

### Fiber Knob Protein Crystallisation

All crystal plates used the Pact Premier Screen (Molecular Dimensions, Suffolk, UK). EasyXtal 15-well Tool crystal plates (Qiagen, Manchester, UK) were set up manually for HAdV-D15, D29 and D30 using the hanging drop method. Drops at either ratios of 1:1 or 3:1 (protein to screen) were suspended over a 300μL reservoir. Crystal plates for HAdV-D24, D25, D32 and D53 fiber-knob proteins were set up using a Mosquito crystallisation robot (SPT Labtech, Hertfordshire, UK) for preliminary work, while optimisation and sialic acid soaking experiments were also performed using manually set up plates.

Plates were sealed and incubated at 18°C for between 7 and 21 days. Crystals of HAdV-D25, D29, D30 and D53 were soaked before harvest with N-Acetyl Neuraminic Acid (Sigma-Aldrich, Gillingham, UK) at a concentration of 10mM.

Crystals were harvested on litholoops (Molecular Dimensions, Suffolk, UK), flash frozen in liquid nitrogen, and transported to the Diamond Light Source where diffraction data were collected on Beamline I03 for HAdV-D15, D25, D29 and D30 and Beamline I04 for HAdV-D24, D25 (with and without sialic acid), D29 soaked with sialic acid and D30 soaked with sialic acid. Data for HAdV-D32 and D53 (with and without sialic acid) were collected on Beamlines I03, I04 and I04-1. During data collection, crystals were maintained at 100°K.

### Structure Determination

Diffraction patterns were recorded and reduced automatically through the Diamond Light Source pipeline with XDS (*21*), xia2 DIALS and 3dii (*22*). Scaling and merging data was completed with Pointless, Aimless and Truncate while MolREP and Phaser were used for structure solving. All of these programs are included in the CCP4 Program package (*23*). RefMac5 was used for map generation and refinement and WinCoot was used for sequence adjustments and model corrections. PyMOL Version 2.2.3 was used to generate illustrations of crystal structures and models (*24*).

### Fiber-Knob Protein Sequence Alignment

Sequences for species D fiber-knob proteins were obtained from NCBI and aligned using Clustal Omega (*25*). The output was displayed in BioEdit.

### Cell Culture

Cells were obtained from the ATCC or collaborators. Cells were passaged twice weekly to maintain a confluency at less than 70% until used in an assay. All cells were cultured at 37°C with 5% CO_2_. SKOV3 cells were cultured in DMEM (Sigma-Aldrich, Gillingham, UK), A549 cells were cultured in RPMI 1640 (Sigma-Aldrich, Gillingham, UK) and BT-20 cells were cultured in α-MEM (Gibco, ThermoFisher Scientific, Waltham, USA). All media was supplemented with 10% fetal bovine serum and 2% penicillin and streptomycin. RPMI 1640 was also supplemented with 1% L-glutamine.

### Neuraminidase Assay

Human cancer cell lines SKOV3, A549 and BT-20 were seeded at 5×10^4^ cells per well in Nunclon Declat Surface 96 well flat-bottomed cell culture plates (Thermo-Fisher Scientific, Waltham, USA). Cells were left to adhere overnight under normal cell culture conditions. Cells were washed with 200μL PBS before neuraminidase (Sigma-Aldrich, Gillingham, UK) was added at 50mU/mL for 1 hour at 37°C. Wells were washed with 200μL PBS before 100μL of replication deficient, luciferase expressing pseudotyped virus in serum free media was added at a viral particle titer of 5,000 VP/cell in triplicate. Plates were incubated for 1 hour on ice. Media containing unbound virus was then discarded, and replaced with 200μL of complete media and plates were incubated at 37°C with 5% CO_2_ for an additional 45 hours. Cells were then washed with 200μL PBS. 5X reporter lysis buffer (Promega, Southampton, UK) was diluted in sterile ddH_2_O and 100μL was added per well. Plates were then frozen at −80°C. Samples were thawed and 10μL of lysate per well was used for a Bicinchoninic Acid Assay (BCA assay). A standard curve of Bovine Serum Albumin (BSA) standards at concentrations 0.025, 0.125, 0.25, 0.5, 0.75, 1.0, 1.5 and 2.0mg/mL was created. Lysate and standards were incubated for 1 hour at 37°C with 5% CO_2_ after the addition of 100μL working reagent (ThermoFisher Scientific, Waltham, USA). Absorbance was measured at 570nm on a plate reader (BioRad, Watford, UK).

To assay for luciferase activity, 100μL luciferase assay substrate (Promega, Southampton, UK) was added to 20μL of the thawed cell lysate in white 96 well plates. Luminescence was quantified using a Clariostar Plus plate reader (BMG, Birmingham, UK). Luminescence was normalised against protein concentration, to give a final value in relative light units (RLU) per milligram (mg) or microgram (μg) of protein.

### Evaluation of CAR binding by HAdV-D fiber knob proteins

CHO cells stably expressing CAR (CHO-CAR) cells were seeded at 2×10^4^ cells per well in 96 well V-bottomed plates. Diluted recombinant fiber knob proteins corresponding to species D adenoviruses at varying concentrations were applied to the appropriate wells. The cells and fiber knob proteins were incubated for 1 hour at 4°C. Wells were made up to 200μL volume and cells were pelleted at 1400 RPM for 4 minutes. The supernatant was removed and 200μL PBS was added to each well. Cells were pelleted again at 1400 RPM for 4 minutes and the supernatant discarded. 50μL of primary antibody (05-644 Anti-CAR clone RmcB, Lot 3022897, EMD Millipore, MA, USA) was added to each well and the cells were incubated on ice at 4°C for 1 hour.

The volume was again made up to 200μL before cells were pelleted at 1400 RPM for 4 minutes. The supernatant was removed, and 200μL PBS added before cells were pelleted using the same process and the supernatant again discarded. 50μL secondary antibody (Goat Anti-Mouse IgG H&L (Alexa Fluor-488), product code: ab150113) was added and the cells were incubated on ice for 30 minutes at 4°C. Cells were again pelleted for 4 minutes by centrifugation at 1400RPM, before being washed with 200uL PBS and pelleted using the same conditions. 100uL PFA was added to each well and the plate was left for 10-15 minutes. 100uL PBS was added on top of the PFA before another centrifugation step. The supernatant was removed and a further wash and centrifugation step performed. Cells were finally resuspended in 200uL PBS and data was collected on an Attune flow cytometer (Invitrogen, ThermoFisher Scientific, Waltham, USA). IC_50_ values were calculated using GraphPad Prism (*26*).

### Statistics

RLU data from the neuraminidase assay was analysed by an unpaired non-parametric Mann-Whitney test in GraphPad Prism (*26*). There were four replicates per condition (n=4). Any results with a p value of less than 0.05 were deemed a statistically significant reduction in infection in the presence of the neuraminidase.

## Results

### High resolution crystal structures confirm conserved structure of species D adenovirus fiber knob proteins

Adenovirus fiber knob proteins crystallised readily, under widely varying pH conditions. The achieved resolution is mostly very high, up to 1.37-1.6Å, although some species gave diffraction to a more modest resolution. Details of data collection and refinement are presented in Supplementary Tables 1-4.

The adenovirus fiber knob protein is known to have a highly conserved trimeric structure. We performed X-ray crystallography to generate structures of species D human adenovirus fiber knob proteins HAdV-D15, D24, D29, D30, D32 and D53 in their apo states. These structures show the expected trimeric form, all viewed down towards the apical domain of the fiber knob proteins (Figure 1).

**Figure 1:**
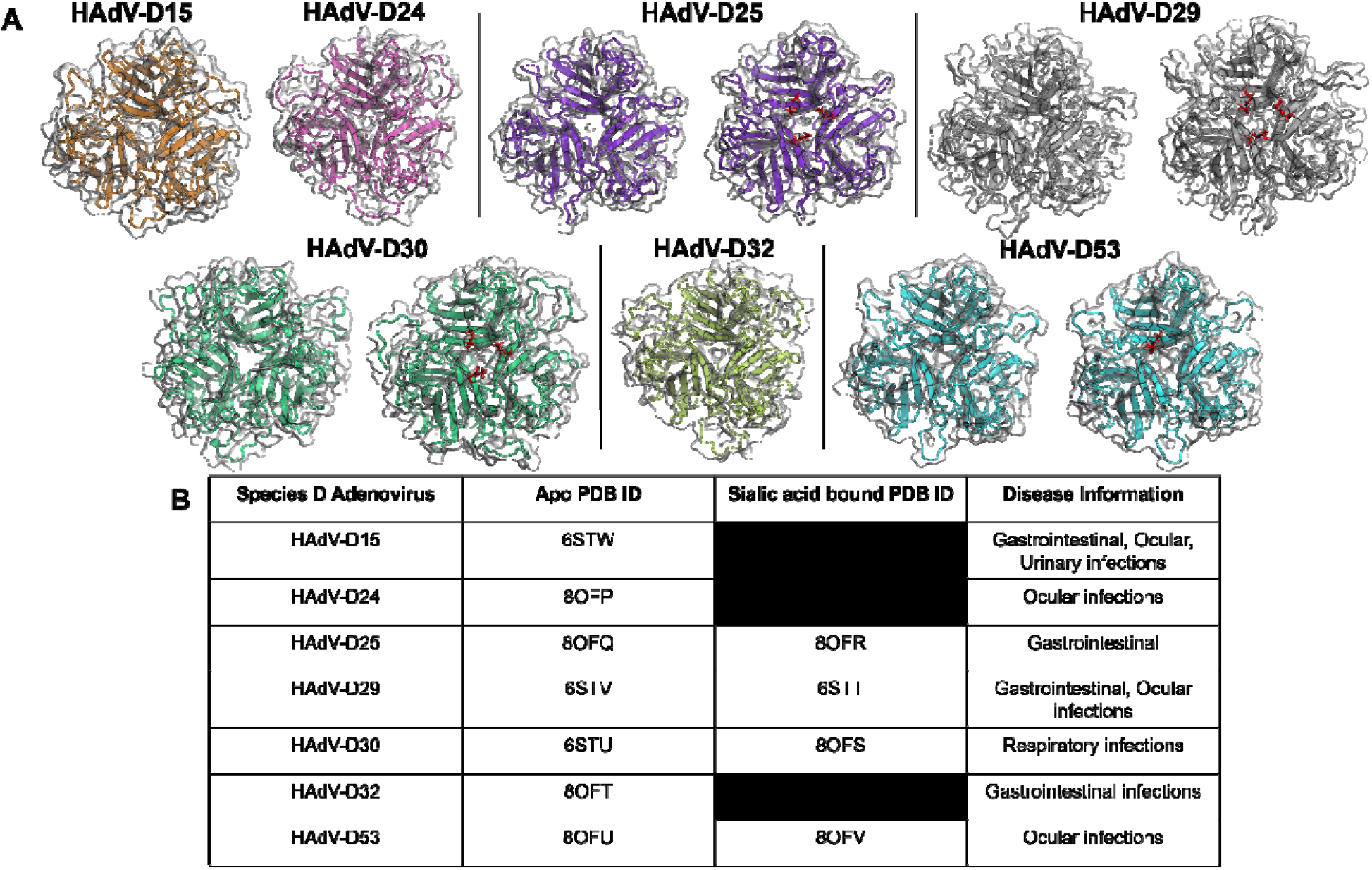
Crystallisation of species D fiber-knob proteins (Charles et al 1995; Shieh 2022; Walsh et al. 2009; Van der Veen and Van der Ploeg 1960; Faden et al. 2005). Adenovirus species D fiber-knob proteins are presented in their apo states in Panel **A**. HAdV-D15, D24 and D32 were crystallised in only their apo states, whereas HAdV-D25, D29, D30 and D53 crystals were also soaked with sialic acid. These structures are shown side by side, with sialic acid highlighted in red. Panel **B** presents the PDB IDs and relevant disease information for each of the species D fiber-knob proteins crystallised.

To evaluate the potential for these recombinant fiber knob proteins to bind cellular sialic acid residues, crystals of the fiber-knob proteins were also soaked with sialic acid prior to data collection. This resulted in four structures of HAdV-D25, D29, D30 and D53 in complex with sialic acid also shown in Figure 1. Sialic acid was observed to bind within a conserved apical region, as observed previously in structures solved for sialic acid interacting species D adenoviruses, including HAdV-D26 (PDB ID: 6QU8) and HAdV-D37 (PDB ID: 1UXA).

The observed space groups were monoclinic (P2_1_ P2 or C2), orthorhombic (P2_1_2_1_2_1_), trigonal (H3 – R3 in hexagonal axes) or cubic (P23). However, there were varying numbers of monomers or trimers in each asymmetric unit, presenting a diverse picture of the fiber knob interactions. HAdV-D24 fiber knob crystals achieved only marginal resolution, and soaking rendered them too disordered for diffraction. Additionally, crystals of HAdV-D32 fiber knob protein soaked with sialic acid could not be obtained.

The conformation of the bound sialic acid was found to be a mixture of the *a* and ß anomers, freely exchangeable in solution and minimal protein side chain adjustment was required to accommodate either conformation. Soaking in sialic acid containing solutions reduced resolution slightly. Crystals of HAdV-D53 fiber knob protein were too sensitive to soaking, and only very short soaks could rescue any samples suitable for diffraction. The short soak allowed only two of four trimers in the asymmetric unit to bind β-sialic acid, with a significant change in packing, explaining the sensitivity of these crystals to soaking.

These adenoviruses are of significant clinical relevance (Figure 1B). They cause gastrointestinal and ocular infections of varying severity. HAdV-D53 is of particular interest due to the recent confirmation of EKC causing infection, similar to previously identified sialic acid binding viruses HAdV-D37, D64 and D19p.

### Conserved residues mark the sialic acid binding pocket

A consensus binding pocket, including bridging waters, was identified in the HAdV-D26 fiber knob protein in complex with sialic acid (PDB ID: 6QU8) anchoring sialic acid in place (*12*). The new structure determinations could not identify the same number of contacts throughout, due to the inability to locate water molecules at different resolutions. Figure 2 panels A-E depicts these pockets in fiber knob proteins of HAdV-D25, D29, D30 and D53, where sialic acid is coloured according to structure and the residues contacting sialic acid are shown in green. Other pocket residues not making contact with sialic acid are shown in red. Figure 2F shows an alignment of the sialic acid binding fiber knob proteins, highlighting contacting residues in yellow compared to the binding pocket identified in HAdV-D26. Y314 and K349 (according to HAdV-D26 sequence) are conserved across all sequences, while the remaining residues are similar, suggesting the existence of a common sialic acid binding pocket across all species fiber knob proteins. Y237 and K349 have also been previously seen to interact in a structure of HAdV-D37 fiber knob protein interacting with sialylactose (*18*).

**Figure 2:**
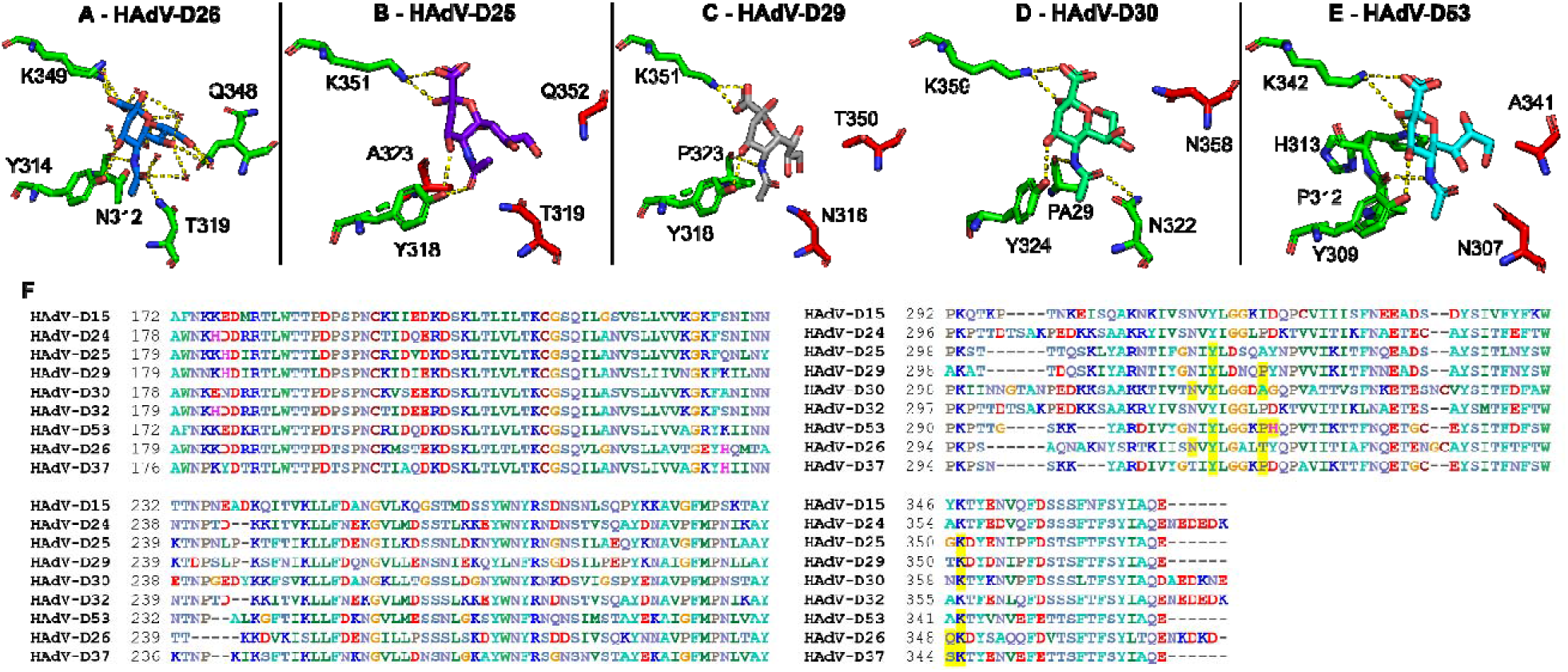
Species D adenoviruses have conserved sialic acid binding residues. Panel **A** shows the sialic acid binding pocket of HAdV-D26 (PDB ID: 6QU8). For all structures, interacting residues are highlighted in green with interactions represented in yellow dashed lines. Residues that are not interacting with sialic acid are highlighted in red. Nitrogen elements are shown in blue, with oxygen elements in red. Sialic acid soaked structures are shown for HAdV-D25 (**B**), D29 (**C**), D30 (**D**) and D53 (**E**). Panel **F** presents an alignment of the fiber-knob region of species D fiber-knob proteins coloured according to residue. Residues highlighted in yellow are shown to be interacting with sialic acid by crystal structure (PDB ID: 1UXA for HAdV-D37 interactions).

### Removal of cell surface sialic acid with neuraminidase reduces the transduction of pseudotyped species D adenoviruses

Following structural investigation of species D adenovirus fiber-knob proteins, *in vitro* cell based assays were used to biologically validate the importance of the identified interactions with sialic acid. Pseudotyped viruses comprising an HAdV-C5 core viral vector psedutoyped with the fiber knob proteins of several HAdV-D of interest (D26, D15, D24, D29 and D53) were generated expressing a luciferase transgene under the control of a ubiquitous CMV IE promoter for measuring transduction following cell treatment with and without neuraminidase treatment.

We used a control, HAdV-C5 expressing luciferase, which has a well characterised and high affinity interaction with CAR that is not influenced by sialic acids (*27*). Alterations in transduction mediated by HAdV-C5 were non-significant as observed across all three cell lines, with a small increase in transduction observed in CAR^LOW^ BT-20 cells (Figure 3A). In contrast, viruses pseudotyped with the fiber-knob protein of HAdV-D26 demonstrated a statistically significant reduction in transduction in all cell lines following cleavage of sialic acid (Figure 3B). This was also demonstrated by viruses pseudotyped with an HAdV-D53 fiber knob protein (Figure 3F), where all reductions in transduction were also statistically significant in all three cell lines tested.

**Figure 3:**
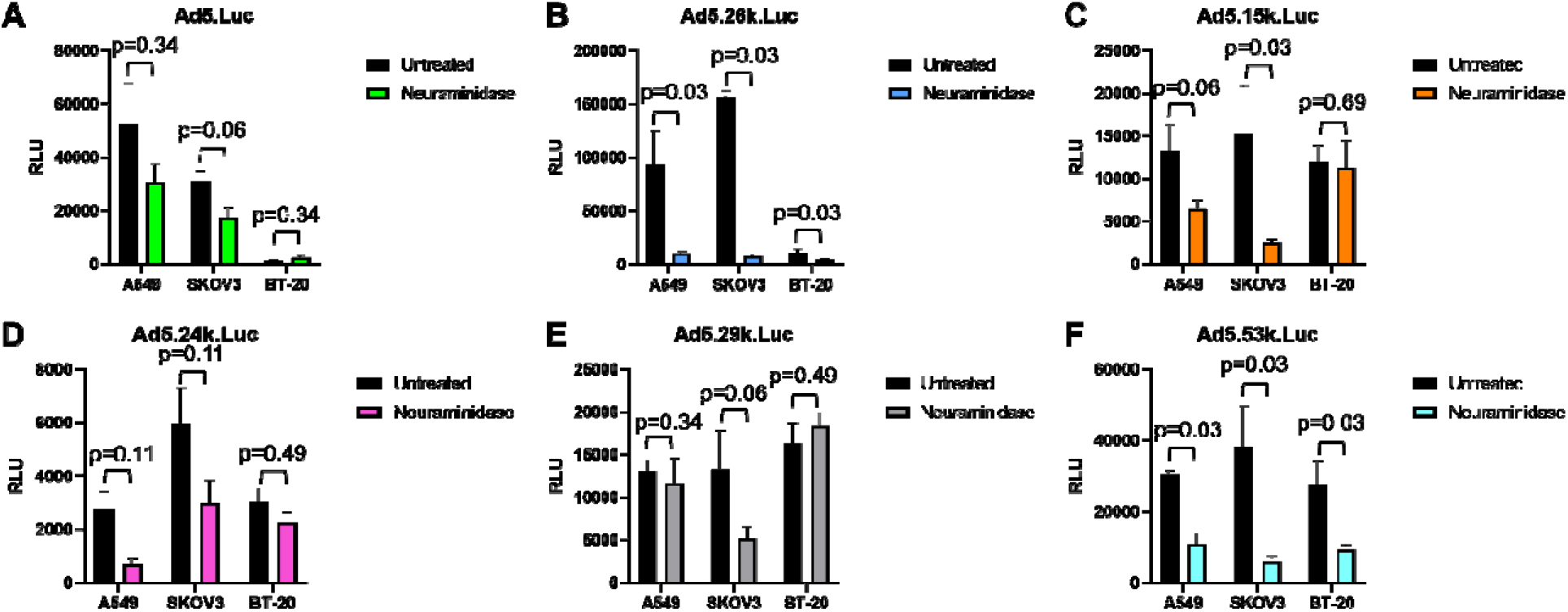
Treatment by neuraminidase reduces transduction of pseudotype species D adenoviruses. A549, SKOV3 and BT-20 cell lines were each infected with luciferase-expressing adenovirus at 5000VP/cell. Panel **A** shows non-significant interaction with HAdV-C5 as a negative control, whereas panel **B** shows statistically significant reduction in transduction by pseudotype HAdV-C5 with a HAdV-D29 fiber-knob protein across all three cell lines. Panels **C, D, E** and **F** present the result of neuraminidase treatment by HAdV-C5 with HAdV-D15, D24, D29 and D53 fiber-knob proteins respectively. Error bars present standard error, and data was statistically tested by Mann-Whitney test (n=4).

Non-significant changes in transduction were observed when cells were transduced with viruses with the fiber-knob proteins of HAdV-D15, D24 and D29 (Figure 3C-E). There was an overall trend in reduction, although in BT-20 cells there was a small, non-significant, increase in transduction of HAdV-D29 fiber-knob protein virus following treatment with neuraminidase. These results indicate significant usage of sialic acid by HAdV-D53 fiber-knob proteins, and the potential partial usage across the other species D adenovirus fiber-knob proteins.

### Characterisation of species D adenovirus fiber knob proteins interactions with CAR

Since CAR is also known to be partially engaged by knob proteins of adenoviruses from HAdV-D including HAdV-D10, D26 and D48 (*13, 28*), we additionally performed binding assays to investigate interactions of species D adenoviruses with CAR (Figure 4). The classical CAR binding adenovirus fiber-knob protein, HAdV-C5, binds CAR with high affinity, with a measured value of 0.0001804μg/10e5 cells. CAR binding was investigated for fiber knob proteins of HAdV-D24, D26 and D32, and was found to be significantly weaker interacting. The respective IC_50_ values were quantified as 0.02931, 0.001261 and 0.01996μg/10e5 cells. These values represent fold changes of 162, 6 and 110 times weaker compared with the IC_50_ value measured for HAdV-C5 fiber knob protein.

**Figure 4:**
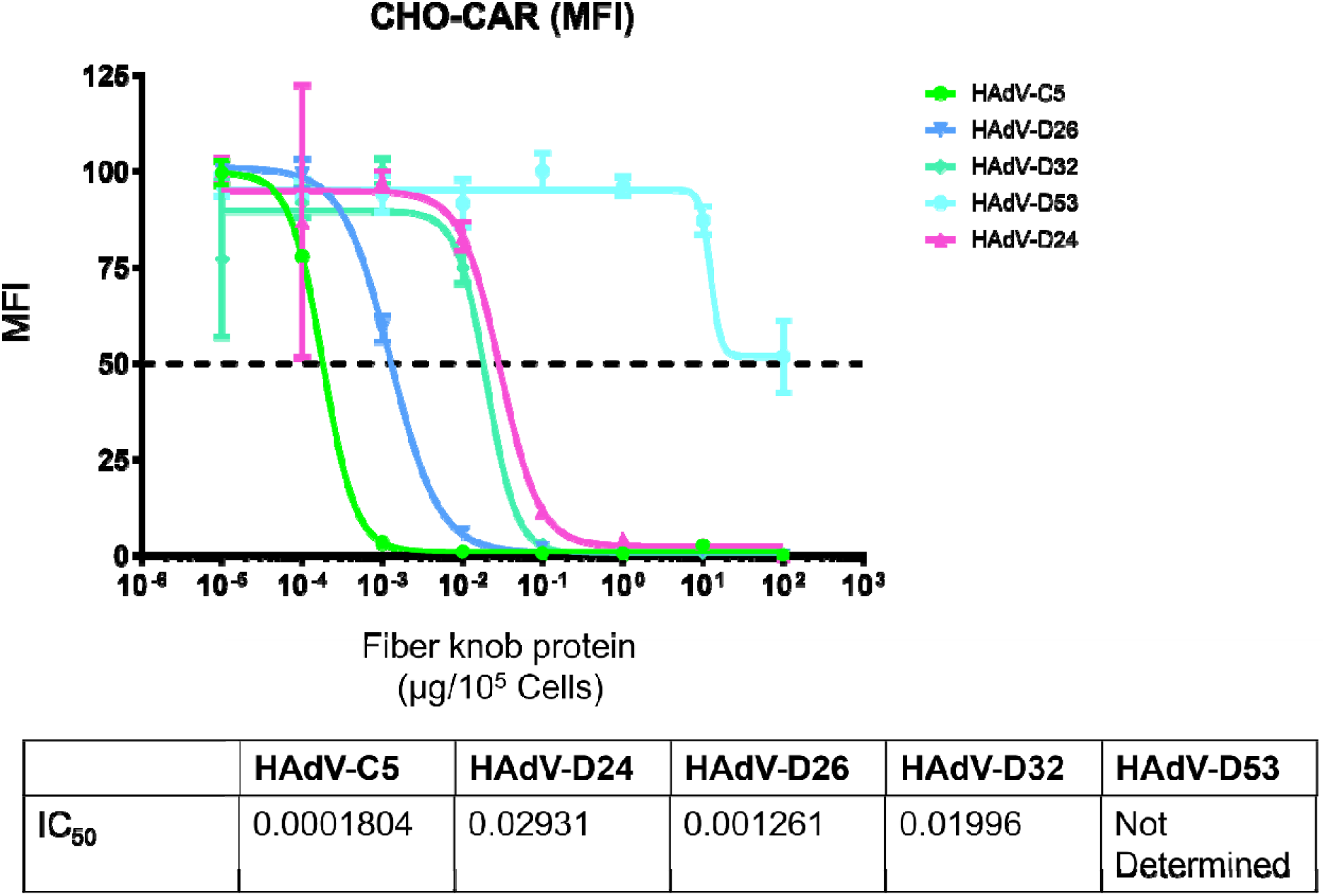
IC_50_ curves suggest varied usage of CAR by species D adenoviruses. Using CHO-CAR cells, adenovirus fiber-knob proteins were tested for interaction with CAR and HAdV-C5 had the strongest IC_50_ value as the positive control. HAdV-D24, D26 and D32 fiber-knob proteins had weaker IC_50_ values but could still interact with CAR. HAdV-D53 fiber-knob protein could not give a reliable IC_50_ value.

The fiber knob protein of HAdV-D53 was also investigated for its ability to interact with CAR. An accurate IC_50_ value could not be measured due to a lack of ability to block CAR antibody binding, even at the highest doses tested. This data suggests variable usage of CAR across the species D adenoviruses’ fiber-knob proteins with some capable of interacting more strongly with CAR than others.

### Inability of HAdV-C5 to bind sialic acid despite apparently conserved residues in the binding pocket

To confirm the validity of our structural observations, we used as a negative control of our structural modelling. We used the fiber-knob protein of HAdV-C5 (PDB ID:6HCN) superposed on the binding pocket of HAdV-D26 (PDB ID: 6QU8) (Figure 5). We observed conservation of binding residues, with the HAdV-C5 chain shown (red) and HAdV-D26 (green) and interactions between sialic acid and HAdV-C5 shown (yellow). Y314 in HAdV-D26, identified in Figure 2 as being universally conserved in species D adenovirus fiber-knob proteins is conserved while K349 is replaced by a histidine in HAdV-C5, a much shorter side chain. This fails to contact sialic acid, weakening the sialic acid pocket which, alongside other substitutions of amino acids shown in Figure 5, makes HAdV-C5 unable to engage sialic acid as species D adenoviruses can.

**Figure 5:**
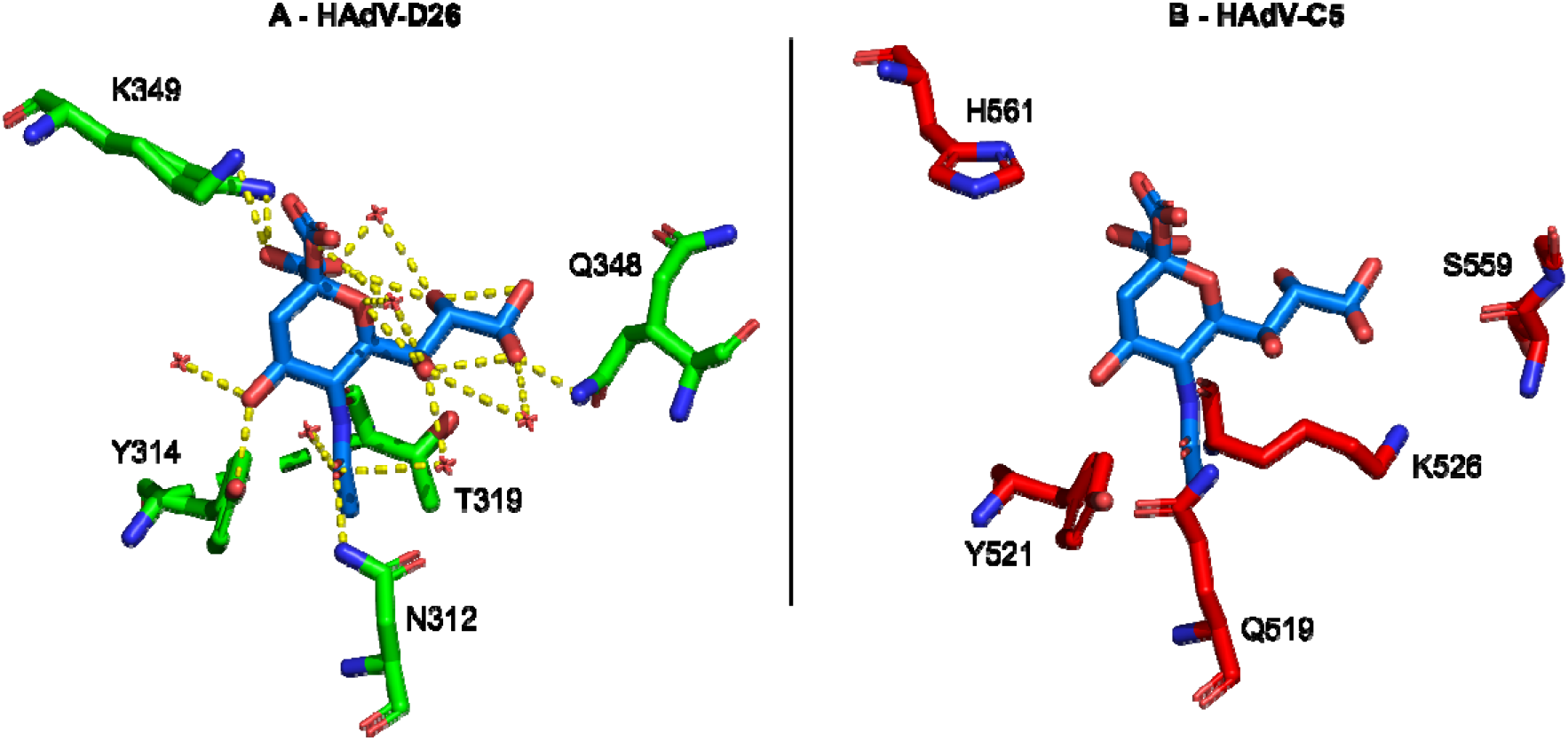
HAdV-C5 fiber-knob protein is incompatible with sialic acid interaction. In panel **A**, The sialic acid binding pocket of HAdV-D26 (PDB ID: 6QU8, shown in green) was aligned to the fiber-knob protein of HAdV-C5 (PDB ID: 6HCN, shown in red) in panel **B** and important residues highlighted. Nitrogen elements are shown in blue with oxygen in red. Many of the residues important in HAdV-D26’s interaction with sialic acid are not present in HAdV-C5, such as the anchoring lysine, preventing interaction with sialic acid.

There is also replacement of T319 with K526 in HAdV-C5, which extends away from the sialic acid in the binding pocket. Whilst the threonine was not found to be universally conserved (Figure 2F), the other species D adenoviruses had similar sized residues which generally made interactions with sialic acid in HAdV-D29, D30 and D53 (Figure 2C, D and E). This contrasts with the lysine seen in the same position in HAdV-C5 which prevents HAdV-C5 binding to sialic acid through steric hindrance. Overall, the residues in the binding pocket in HAdV-C5 are shown to differ from those seen in the binding pockets in Figure 2, highlighting key differences between sialic acid utilising viruses, and those known to use different receptors such as CAR. This suggests that the sialic acid binding pocket identified in species D adenovirus fiber-knob proteins is not universal across all species and may be a unique feature of HAdV-D.

## Discussion

Sialic acid is well characterised as a primary cell receptor for a wide range of viruses including influenza and some coronaviruses, in addition to human adenoviruses (*29*). Previous studies have identified sialic acid interactions with individual species D viruses including HAdV-D26, 37 and 64 (*12, 17, 19*). Low sequence similarity in the fiber-knob protein has been seen to be tolerated to still allow usage of sialic acid cell entry (*12*), hinting at broader usage of sialic acid across HAdV-D. Although there is clear sequence variation across the proteins, we observed conservation of key sialic acid binding residues as highlighted in the alignment and the individual crystal structure for each complexed protein (Figure 2A-E). The fiber-knob protein initiates cell entry by human adenoviruses and so binding here is critical for infection. These interactions are pivotal in defining a viral tropism, and knowledge of these interactions can aid in design of antivirals for therapeutic purposes. It should be noted that another adenovirus from a different species has also been identified as binding sialic acid. Species G virus HAdV-G52 fiber-knob protein binds sialic acid, however, it does so at a different binding site on the side of the fiber-knob protein (*30*). This further suggests the apical binding pocket may be specific to the species D adenoviruses, whilst alternative means to engage sialic acids may be conserved across other species.

Previous studies have identified an ocular tropism in humans common among sialic acid utilising viruses (*31*). The species D adenoviruses generally cause rare gastrointestinal or ocular infections (Figure 1B). Many previously identified species D viruses that utilise sialic acid cause EKC, the most severe form of human adenovirus caused conjunctivitis (*32*). Particularly contagious, EKC is characterised by its involvement of the cornea, can result in blindness and is notoriously difficult to treat with the potential to cause chronic disease (*33*). One of the viruses in this article that demonstrated significantly reduced transduction on removal of sialic acid following neuraminidase treatment is HAdV-D53 (Figure 3F). This virus has been confirmed to also cause EKC and has been isolated from patients in Germany and Japan (*34, 35*). The species D adenoviruses are notable for their frequent recombination events and when investigated, HAdV-D53 seemed to have arisen from HAdV-D22 recombining with two sialic acid binding, EKC causing adenoviruses: HAdV-D8 and D37 (*34, 35*). This recombination event, combining two viruses with similar tropisms explains our findings that HAdV-D53 appears to rely heavily on sialic acid as a cell entry receptor (Figure 2E and Figure 3F), and draws another similarity between the EKC causing adenoviruses.

The variation in sialic acid and CAR usage seen across the species D viruses investigated in this article suggests that the viruses within this species may not rely on a single receptor for initiating cell infection. The species C viruses have well-defined primary cell entry receptor interactions with CAR (*27*). The interactions between HAdV-C5 and CAR is especially well characterised, as HAdV-C5 forms the basis of many vectors in previous and current clinical trials including as a vaccine platform for COVID-19 and as a vector in a newly approved therapy for bladder cancer (*36, 37*). In contrast, HAdV-B viruses display dual receptor usage similar to our observations of the species D adenoviruses in this article and are known to use both CD46 and in some cases, desmoglein-2 (DSG-2) (*38, 39*). It is clear from the data in Figure 3 that a single species wide receptor is unlikely for HAdV-D. This led us to investigate whether dual receptor usage was employed by HAdV-D.

The classical human adenovirus receptor, CAR, is over-expressed on CHO-CAR cells used for our binding experiments (Figure 4) which demonstrated variable affinity of species D fiber-knob proteins for CAR. As expected, this interaction was much weaker than that of HAdV-C5 fiber-knob protein. Fiber-knob proteins of HAdV-D24, D26 and D32 all had a much weaker IC_50_ value, with HAdV-D53’s fiber-knob protein seemingly unable to interact with CAR. This again suggested a range of variability, with some viruses such as HAdV-D53 having a clear preference for sialic acid primarily but others demonstrating the ability to interact with both receptors.

Sialic acids are a family of terminal monosaccharides at the end of the glycan chains that decorate many vertebrate cells, making it a highly biologically relevant receptor but also a receptor of variable nature making it difficult to investigate (*20*). This means we must also consider the possibility that variation in glycan chains could change the ability for usage by species D adenoviruses. Few viruses are known to have a glycan chain preference, with HAdV-D37 known to preferentially bind GD1a glycan (*40*) and so it is important to consider this when interpreting our results as different fiber-knob proteins may have a preference for glycosylation patterns which may affect their usage of sialic acid and CAR. This is all before considering secondary interactions with the hexon and penton proteins of the adenovirus capsid demonstrating the complexity of this interaction and the difficulty in defining it.

Soaking our fiber-knob protein crystals allowed direct investigation into the potential for sialic acid to interact in the previously identified binding pocket. In contrast, modelling this interaction *in silico* could have reduced the accuracy that comes with X-ray crystallography. In this article we present seven novel structures of apo state fiber-knob proteins of species D adenoviruses (Figure 1A). These provide a good basis from which to conduct structural analyses. However, the four sialic acid soaked novel crystal structures (Figure 2B-E), provide a much more reliable view of the binding pocket and interactions with sialic acid. However, it is important to remember that crystallography shows only a snapshot of the interaction, and not a dynamic model. This limitation encouraged the usage of an integrated structural biological workflow, to couple our structural observations with biologically measured effects.

By studying a virus’ protein structure we can begin to elucidate it’s basic mechanisms of cellular infection. This can confer benefits in two main applications. Primarily, understanding such mechanisms is vitally important when designing antivirals to combat the virus and the diseases they cause. In the case of EKC causing viruses, infection can result in chronic disease costing healthcare providers huge amounts over patients’ lifetimes and drastically reducing patients’ quality of life (*32*). The second application considers the therapeutic potential of species D adenoviruses. Traditionally HAdV-C5 has been the basis of many adenovirus derived vectors, however, the virus has high seroprevalence in many populations resulting in high levels of neutralising antibodies (*41*). The rarity of infections by species D adenoviruses means there is low pre-existing immunity to the viruses and so they could be effective alternative platforms to HAdV-C5. There is already some precedence to using species D adenoviruses in therapeutics, with HAdV-D26 being used as a vaccine platform to combat Ebola, HIV and SARS-CoV2 infections (*42–44*). We can only achieve these two benefits if we have a solid basis of understanding of these viruses which is where studies investigating structure and function can help to build a profile of each individual virus, to ameliorate its burden on both patient and healthcare system but also to utilise the virus to our advantage in a therapeutic setting.

Future work in this area should focus on defining the preference of the individual species D adenoviruses for each receptor. This article highlights the variation across this species, unlike the highly conserved tropisms seen across species B and C viruses (*11, 38*), and therefore it is likely that each individual virus will need to be profiled prior to the design of antivirals or usage as a therapeutic vector. Previous studies have highlighted the species D viruses’ rarity of infection (*45*) and therefore their lack of neutralising antibodies present in patients. Thus, they represent a promising option for future therapeutics, but must be properly vetted prior to being used as vectors.

Our findings demonstrate a conserved binding site for sialic acid, common across the species D adenoviruses, therefore suggesting that, along with CAR, the receptor can be used to initiate cell entry. When comparing to previously known sialic acid binders, we can appreciate how the diversity of sequence across the species D fiber-knob proteins is tolerated and still permits interaction with sialic acid. We validated our structural data with *in vitro* assays which confirmed species D fiber-knob proteins were capable of interacting with both CAR and sialic acid, raising the possibility of variable usage of both receptors across the species. This will be important when designing antiviral drugs for the species D viruses, especially those such as HAdV-D37 and D53 which have the capability to cause EKC creating a large burden on healthcare systems. On a wider basis, the species D viruses have great potential as therapeutic vectors but more must be done to understand their mechanisms of infection prior to them being used in the clinic.

## Supporting information

Supplementary Figures and Tables, Mundy RM et al.

## Acknowledgments

The authors would like to thank Diamond Light Source for beamtime (proposals mx18812, mx20147 and mx29990), and the staff of beamlines I03, I04 and I04-1 for assistance with crystal testing and data collection.

## Author contributions

Conceptualization: RMM, ATB, PJR, ALP

Methodology: RMM, ATB, EAB, EM, PJR, ALP

Formal analysis: RMM, ATB, PRJ, ALP

Investigation: RMM, ATB, EAB, EM, PJR, TGC, AT-C

Visualization: RMM, EAB, PJR

Supervision: PJR, ALP

Writing—original draft: RMM, PJR, ALP

Writing—review & editing: RMM, ATB, EAB, EM, PJR, ALP, TGC, AT-C Funding acquisition: ALP

## Data and materials availability

Material described in this manuscript can be made available under a material transfer agreement (MTA). Accession numbers (PDB) for the structures outlined are included in the text. All data are available in the main text or the supplementary materials.

