## Supplementary Figures and Tables, Mundy RM et al. for "Broad sialic acid usage amongst species D human adenovirus"

Rosie M. Mundy *et al.*

**This PDF file includes:**

Supplementary Text

Figs. S1

Tables S1 to S4

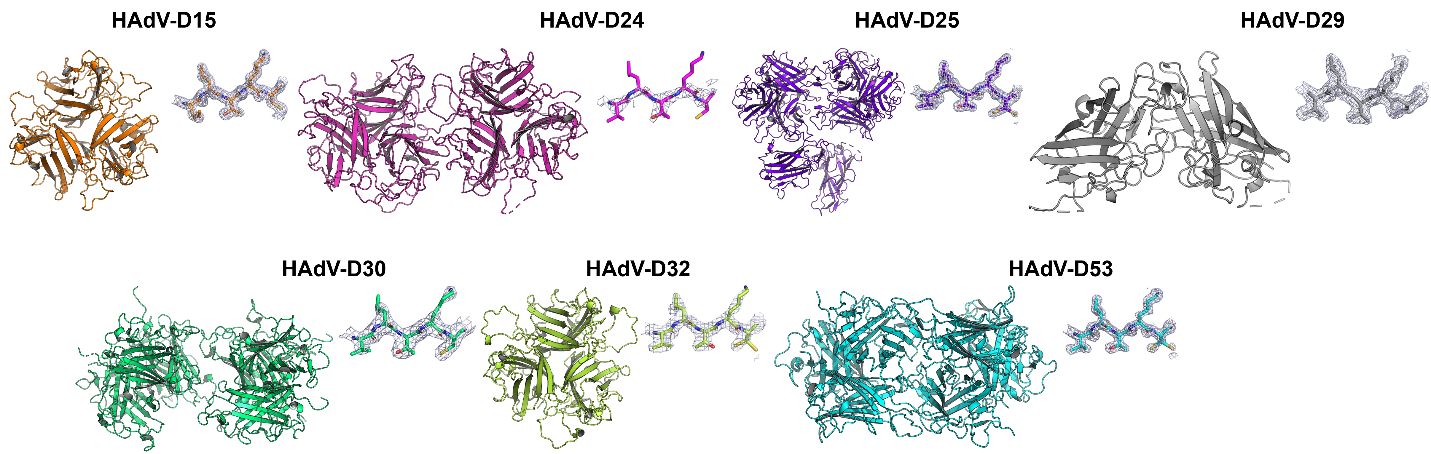

**Figure S1.: Asymmetric units of apo state structures of novel adenovirus fiber-knob proteins.** Each fiber-knob protein structure is depicted in it’s apo state, with an example region of the density map shown adjacent to each unit. The fiber-knob proteins are coloured according to the same code as in the main figures.

Table S1. Data collection and refinement statistics for HAdV-D24 and D25 fiber-knob protein X-ray crystallography structures generated.

| **Protein** | **HAdV-D25 Apo State** | **HAdV-D25 Sialic Acid Soak** | **HAdV-D24 Apo State** |
| --- | --- | --- | --- |
| **PDB Entry** | **8OFQ** | **8OFR** | **8OFP** |
| **Data Collection** | | | |
| Diamond Beamline | DLS_I04 | DLS_I04 | DLS_I04 |
| Date | 2019-10-17 | 2019-07-15 | 2019-10-17 |
| Wavelength | 0.9795 | 0.88561 | 0.9795 |
| Crystallisation Conditions | 0.1 M SPG buffer, 25 % w/v PEG 1500 | 0.1 M SPG buffer, 25 % w/v PEG 1500 | 0.1 M MMT Buffer, 25% PEG 1500 |
| pH | 6.0 | 6.0 | 5.0 |
| **Crystal Data** | | | |
| *a,b,c* (Å) | 126.55, 126.55, 251.44 | 127.56, 179.32, 207.9 | 51.259, 99.619, 113.55 |
| a,b,g (°) | 90.0, 90.0, 120.0 | 90.0, 90.0, 90.0 | 90.0, 100.26, 90.0 |
| Space group | H3 ( R3 hexagonal axes) | P 21 21 21 | P 1 21 1 |
| Resolution (Å) | 1.599 – 82.62 | 2.51 – 64.65 | 2.90 - 55.87 |
| Outer shell | 1.60 – 1.64 | 2.51 – 2.58 | 2.90 – 2.98 |
| *R-*merge (%) | 7.1 (128.4 | 21.8 (226.8) | 15.0 (111.7) |
| *R-*pim (%) | 4.5 (81.5) | 11.7 (120.7) | 8.9 (64.9) |
| *R-*meas (%) | 7.8 (141.8) | 23.4 (242.7) | 17.5 (129.6) |
| CC1/2 | 0.999 (0.484) | 0.988 (0.451) | 0.973 (0.652) |
| I / σ(I) | 11.2 (1.1) | 6.0 (0.9) | 4.4 (1.0) |
| Completeness (%) | 100.0 (100.0) | 100.0 (100.0) | 100.0 (100.0) |
| Multiplicity | 5.8 (5.6) | 7.7 (7.8) | 3.8 (3.9) |
| Total Measurements | 1,144,577 (82,532) | 1,247,870 (93,395) | 96,302 (7,260) |
| Unique Reflections | 198,233 (14,640) | 162,833 (11,926) | 25,128 (1,852) |
| Wilson B-factor(Å^2^) | 21.7 | 48.9 | 67.3 |
| Monomers / au | 8 | 24 | 6 |
| Trimer / au | 2 + 1/3 + 1/3 | 8 | 2 |
| **Refinement Statistics** | | | |
| Resolution range | 1.60 – 82.61 | 2.51 – 64.64 | 3.10 – 55.87 |
| Non-H Atoms | 12,828 | 36,081 | 8946 |
| R-work reflections | 188,438 | 154,712 | 19,502 |
| R-free reflections | 9,768 | 7,943 | 1,004 |
| R-work/R-free (%) | 16.8 / 20.3 | 22.0 / 23.9 | 23.4 / 33.7 |
| **^2^rms deviations** | | | |
| Bond lengths (Å) | 0.013 | 0.010 | 0.009 |
| Bond Angles (°) | 1.924 | 1.295 | 1.371 |
| ^3^Coordinate error | 0.070 | 0.307 | 0.72 |
| Mean B value (Å^2^) | 28.0 | 56.5 | 94.0 |
| **Ramachandran Statistics** | | | |
| Favoured/allowed/Outliers | 1196 / 59 / 8 | 3579 / 262 / 41 | 833 / 163 / 111 |
| % | 94.7 / 4.7 / 0.6 | 92.2 / 6.8 / 1.1 | 75.3 / 14.7 / 10.0 |

* One crystal was used for determining the structure.

* Figures in brackets refer to outer resolution shell, where applicable.

^2^Figures in brackets are rms targets

^3^Coordinate Estimated Standard Uncertainty in (Å), calculated based on maximum likelihood statistics.

Table S2.: Data collection and refinement statistics for HAdV-D15 and D29 fiber-knob protein X-ray crystallography structures generated.

| **Protein** | **HAdV-D15 Apo State** | **HAdV-D29 Apo State** | **HAdV-D29 Sialic Acid Soak** |
| --- | --- | --- | --- |
| **PDB Entry** | **6STW** | **6STV** | **6STT** |
| **Data Collection** | | | |
| Diamond Beamline | I03 | I03 | I04 |
| Date | 18/04/2019 | 18/04/2019 | 18/05/2019 |
| Wavelength | 0.95372 | 0.95372 | 0.9795 |
| Crystallisation Conditions | 0.1M SPG, 25% w/v PEG 1500 | 0.1M SPG, 25% w/v PEG 1500 | 0.1M SPG, 25% w/v PEG 1500 |
| pH | 8.0 | 6.0 | 6.0 |
| **Crystal Data** | | | |
| *a,b,c* (Å) | 59.58, 89.18, 106.20 | 104.29, 104.29, 104.29 | 104.24, 104.24, 104.24 |
| a,b,g (°) | 90, 90, 90 | 90, 90, 90 | 90, 90, 90 |
| Space group | P 2_1_ 2_1_ 2_1_ | P 2 3 | P 2 3 |
| Resolution (Å) | 1.37 – 39.64 | 1.60 – 73.85 | 1.54 – 60.18 |
| Outer shell | 1.37 – 1.41 | 1.60 – 1.63 | 1.54 – 1.57 |
| *R-*merge (%) | 0.036 (1.604) | 0.105 (3.525) | 9.3 (237.0) |
| *R-*pim (%) | 2.1 (93.6) | 2.3 (77.6) | 2.0 (51.2) |
| *R-*meas (%) | 4.1 (186.2) | 10.9 (369.0) | 9.6 (242.5) |
| CC1/2 | 1.000 (0.478) | 1.000 (0.444) | 1.000 (0.605) |
| I / σ(I) | 18.4 (1.1) | 18.1 (1.0) | 17.9 (1.4) |
| Completeness (%) | 100.0 (100.0) | 100.0 (100.0) | 100.0 (100.0) |
| Multiplicity | 7.4 (7.6) | 22.6 (22.5) | 22.4 (22.3) |
| Total Measurements | 881,087 (66,057) | 1,131,680 (54,939) | 1,253,942 (62,276) |
| Unique Reflections | 119,296 (8,721) | 50.042 (2,447) | 55,982 (2,787) |
| Wilson B-factor(Å^2^) | 19.9 | 24.2 | 20.9 |
| Monomers / au | 3 | 2 | 2 |
| Trimer / au | 1 | 1/3 + 1/3 | 1/3 + 1/3 |
| **Refinement Statistics** | | | |
| Resolution range | 1.37 – 39.64 | 1.60 – 73.85 | 1.54 – 60.25 |
| Non-H Atoms | 5,429 | 3,247 | 3,290 |
| R-work reflections | 113,301 | 47,543 | 53,165 |
| R-free reflections | 5,905 | 2,470 | 2,767 |
| R-work/R-free (%) | 17.3 / 19.9 | 17.6 / 20.1 | 14.0 / 18.8 |
| **^2^rms deviations** | | | |
| Bond lengths (Å) | 0.012 | 0.012 | 0.013 (0.013) |
| Bond Angles (°) | 1.722 | 1.699 | 1.668 (1.649) |
| ^3^Coordinate error | 0.049 | 0.075 | 0.061 |
| Mean B value (Å^2^) | 27.8 | 30.0 | 31.0 |
| **Ramachandran Statistics** | | | |
| Favoured/allowed/Outliers | 557 / 33 / 2 | 338 / 16 / 0 | 336 / 13 / 2 |
| % | 94 / 6 / 0 | 96 / 4 / 0 | 96 / 4 / 1 |

* One crystal was used for determining the structure.

* Figures in brackets refer to outer resolution shell, where applicable.

^2^Figures in brackets are rms targets

^3^Coordinate Estimated Standard Uncertainty in (Å), calculated based on maximum likelihood statistics.

Table S3.: Data collection and refinement statistics for HAdV-D30 and D32 fiber-knob protein X-ray crystallography structures generated.

| **Protein** | **HAdV-D30 Apo State** | **HAdV-D30 Sialic Acid Soak** | **HAdV-D32 Apo State** |
| --- | --- | --- | --- |
| **PDB Entry** | **6STU** | **8OFS** | **8OFT** |
| **Data Collection** | | | |
| Diamond Beamline | DLS_I03 | DLS_I04 | DLS_I04-1 |
| Date | 2019-04-18 | 2019-05-18 | 2019-12-14 |
| Wavelength | 0.95372 | 0.9795 | 0.91188 |
| Crystallisation Conditions | 0.1M SPG, 25% w/v PEG 1500 | 0.1M MMT [Malic acid, MES, Tris], 25% w/v PEG 1500 | 0.1 M SPG Buffer, 25% PEG 1500 |
| pH | 6.0 | 8.0 | 6.0 |
| **Crystal Data** | | | |
| *a,b,c* (Å) | 63.35,87.36,217.86 | 62.13, 62.49, 71.8 | 67.043, 93.748, 94.793 |
| α,β,γ (°) | 90, 90, 90 | 90.0, 93.85, 90.0 | 90.0, 98.31, 90.0 |
| Space group | P 2_1_ 2_1_ 2_1_ | P 1 21 1 | C 1 2 1 |
| Resolution (Å) | 2.39 – 54.76 | 2.56 – 62.5 | 2.0 - 93.80 |
| Outer shell | 2.39 – 2.45 | 2.56 – 2.63 | 2.0 – 2.05 |
| *R-*merge (%) | 0.116 (1.376) | 5.7 (97.2) | 10.2 (55.1) |
| *R-*pim (%) | 4.9 (56.1) | 3.4 (57.0) | 6.2 (35.3) |
| *R-*meas (%) | 0.134 (1.583) | 6.6 (113.1) | 11.9 (65.8) |
| CC1/2 | 0.998 (0.623) | 0.998 (0.824) | 0.992 (0.831) |
| I / σ(I) | 11.2 (1.4) | 10.5 (1.1) | 5.9 (1.4) |
| Completeness (%) | 100.0 (100.0) | 99.0 (99.3) | 99.4 (94.8) |
| Multiplicity | 7.5 (7.8) | 3.7 (3.8) | 3.6 (3.3) |
| Total Measurements | 365,880 (28,110) | 66,423 (5,020) | 139,963 (9,081) |
| Unique Reflections | 48,897 (3,589) | 17,735 (1,327) | 39,152 (2,762) |
| Wilson B-factor(Å^2^) | 48.7 | 74.3 | 28.0 |
| Monomers / au |  | 6 | 3 |
| Trimer / au |  | 2 | 1 |
| **Refinement Statistics** | | | |
| Resolution range | 2.39-54.76 | 2.56 – 48.56 | 2.00 – 93.80 |
| Non-H Atoms |  | 4,701 | 5,011 |
| R-work reflections | 46,478 | 16,730 | 36,587 |
| R-free reflections | 2,348 | 933 | 1,834 |
| R-work/R-free (%) | 21.7 / 26.8 | 22.7 / 30.0 | 21.2 / 25.1 |
| **^2^rms deviations** | | | |
| Bond lengths (Å) | 0.010 | 0.006 | 0.012 |
| Bond Angles (°) | 1.722 | 1.537 | 1.772 |
| ^3^Coordinate error | 0.293 | 0.896 | 0.168 |
| Mean B value (Å^2^) | 58.7 | 125.7 | 39.1 |
| **Ramachandran Statistics** | | | |
| Favoured/allowed/Outliers | 1122 / 78 / 8 | 521 / 61 / 13 | 514 / 36 / 10 |
| % | 92.88 / 6.46 / 0.66 | 87.6 / 10.3 / 2.2 | 91.8 / 6.4 / 1.8 |

* One crystal was used for determining the structure.

* Figures in brackets refer to outer resolution shell, where applicable.

^2^Figures in brackets are rms targets

^3^Coordinate Estimated Standard Uncertainty in (Å), calculated based on maximum likelihood statistics.

Table S4.: Data collection and refinement statistics for HAdV-D53 fiber-knob protein X-ray crystallography structures generated.

| **Protein** | **HAdV-D53 Apo State** | **HAdV-D53 Sialic Acid Soak** |
| --- | --- | --- |
| **PDB Entry** | **8OFU** | **8OFV** |
| **Data Collection** | | |
| Diamond Beamline | I04-1 | I04-1 |
| Date | 14 – 12 – 2019 | 14 – 12 - 2019 |
| Wavelength | 0.91188 | 0.91188 |
| Crystallisation Conditions | 0.2 M potassium thiocyanate, 20 % w/v PEG 3350 | 0.1 M PCTP buffer, 25 % w/v PEG 1500 |
| pH | Unadjusted | 4.0 |
| **Crystal Data** | | |
| *a,b,c* (Å) | 206.314, 60.526, 93.837 | 93.55, 60.406, 243.686 |
| a,b,g (°) | 90.0, 91.796, 90.0 | 90.0, 99.88, 90.0 |
| Space group | C 1 2 1 | P 1 2 1 |
| Resolution (Å) | 1.606 – 70.489 | 1.77 - 92.17 |
| Outer shell | 1.606 – 1.633 | 1.77 – 1.80 |
| *R-*merge (%) | 9.9 (156.9) | 16.8 (170.7) |
| *R-*pim (%) | 5.9 (97.2) | 9.7 (96.6) |
| *R-*meas (%) | 11.6 (185.3) | 19.5 (196.7) |
| CC1/2 | 0.998 (0.334) | 0.685 (0.327) |
| I / σ(I) | 8.7 (0.8) | 6.0 (0.6) |
| Completeness (%) | 99.7 (99.4) | 98.7 (97.7) |
| Multiplicity | 3.7 (3.5) | 3.8 (3.9) |
| Total Measurements | 563,343 (26,448) | 981,049 (49,583) |
| Unique Reflections | 150,644 (7,529) | 258,349 (12,686) |
| Wilson B-factor(Å^2^) | 20.6 | 17.4 |
| Monomers / au | 6 | 12 |
| Trimer / au | 2 | 4 |
| **Refinement Statistics** | | |
| Resolution range | 1.61 – 68.42 | 1.77 – 92.17 |
| Non-H Atoms | 10,116 | 19,836 |
| R-work reflections | 142,637 | 245,379 |
| R-free reflections | 7,463 | 12,949 |
| R-work/R-free (%) | 18.2 / 20.8 | 20.8 / 24.5 |
| **^2^rms deviations** | | |
| Bond lengths (Å) | 0.014 (0.013) | 0.011 (0.013) |
| Bond Angles (°) | 1.825 (1.652) | 1.702 (1.660) |
| ^3^Coordinate error | 0.087 | 0.146 |
| Mean B value (Å^2^) | 25.1 | 25.4 |
| **Ramachandran Statistics** | | |
| Favoured/allowed/Outliers | 986 / 47 / 6 | 1927 / 76 / 18 |
| % | 94.9 / 4.5 / 0.6 | 95.4 / 3.8 / 0.9 |

* One crystal was used for determining the structure.

* Figures in brackets refer to outer resolution shell, where applicable.

^2^Figures in brackets are rms targets

^3^Coordinate Estimated Standard Uncertainty in (Å), calculated based on maximum likelihood statistics.
